# Inferring functional organization of posterior parietal cortex circuitry based on information flow

**DOI:** 10.1101/2023.03.08.531516

**Authors:** Jung Uk Kang, Eric Mooshagian, Lawrence H. Snyder

## Abstract

Many studies infer the role of neurons by asking what information can be decoded from their activity or by observing the consequences of perturbing their activity. An alternative approach is to consider information flow between neurons. We applied this approach to the Parietal Reach Region (PRR) and the Lateral Intraparietal area (LIP) in posterior parietal cortex. Two complementary methods show that, across a range of reaching tasks, information flows primarily from PRR to LIP but not vice versa. This suggests that PRR determines the spatial goals of coordinated eye and arm movements and instructs LIP of those goals. Based on these findings, we conclude that PRR and LIP operate in a parallel rather than hierarchical manner to plan arm and eye movements, respectively. Similar methodology can be applied to other areas to infer their relative relationships.

## Introduction

There is a rich history of identifying modules within the brain and assigning roles to them based either on what function(s) are perturbed following an intervention or by what information their neurons encode^1–4^. An alternative approach is to consider information flow, based on anatomical or functional measures^5–10^. Functional connectivity can be correlative or causal (effective connectivity) and may depend on the task being performed, providing additional insight into functional organization^11–16^. In this study, we assayed the direction of information flow between saccade and reach areas in posterior parietal cortex during coordinated eye and arm movements by considering temporal relationships between action potentials (spikes) and local field potentials (LFP) as well as LFP-LFP relationships^16^. By applying directed connectivity measures across a range of eye and arm movement tasks, we were able to resolve a long-standing debate involving the relative roles of posterior parietal areas in spatial processing.

Various posterior parietal areas have been associated with particular effectors and movement types. As examples, the anterior intraparietal area is associated with grasping movements^17–19^, the medial superior temporal area with pursuit eye movements^20, 21^, the parietal reach region (PRR) with reaching movements^22, 23^, and the lateral intraparietal area (LIP) with saccadic eye movements^24–26^. Lesions and electrophysiological data suggest that LIP and PRR may be involved in eye-hand coordination^27–30^. LIP has also been implicated in higher-order cognitive aspects of information processing, including directing attention and assigning value to particular locations in space^31–38^. LIP’s roles in saccades and spatial attention are linked by the fact that gaze shifts are a critical mechanism for directing attention^39–41^. To capture these different roles, LIP is often described as forming a “priority map” of space^35, 36^. This description emphasizes what is coded by LIP. In our view, it is critical to consider what role an area plays in supporting brain processes. We therefore interpret the notion of a priority map as a proposal that LIP serves as a general spatial command center, selecting locations of interest and then distributing those spatial signals to other cognitive and motor areas, appropriate to the task at hand. We set out to test this proposal.

Whether area LIP acts as a general spatial command center or plays a more specific role in saccade planning leads to different predictions about the direction of information flow between PRR and LIP during coordinated eye and arm movements. If LIP plays a command role, then information should flow primarily from LIP to reach planning areas like PRR (Fig. 1a). If instead LIP is part of an oculomotor pathway and not involved in general spatial selection, then the two areas might reciprocally interact. Depending on the task, PRR might even influence LIP regarding where to direct a saccade (Fig. 1b). We focused on the high beta band (15-40 Hz), which has been implicated in motor planning^42, 43^. Our results suggest that PRR determines the spatial goal of coordinated eye and arm movements and that PRR and LIP operate in parallel to process arm and eye movements, respectively (Fig. 1b).

**Fig. 1.**
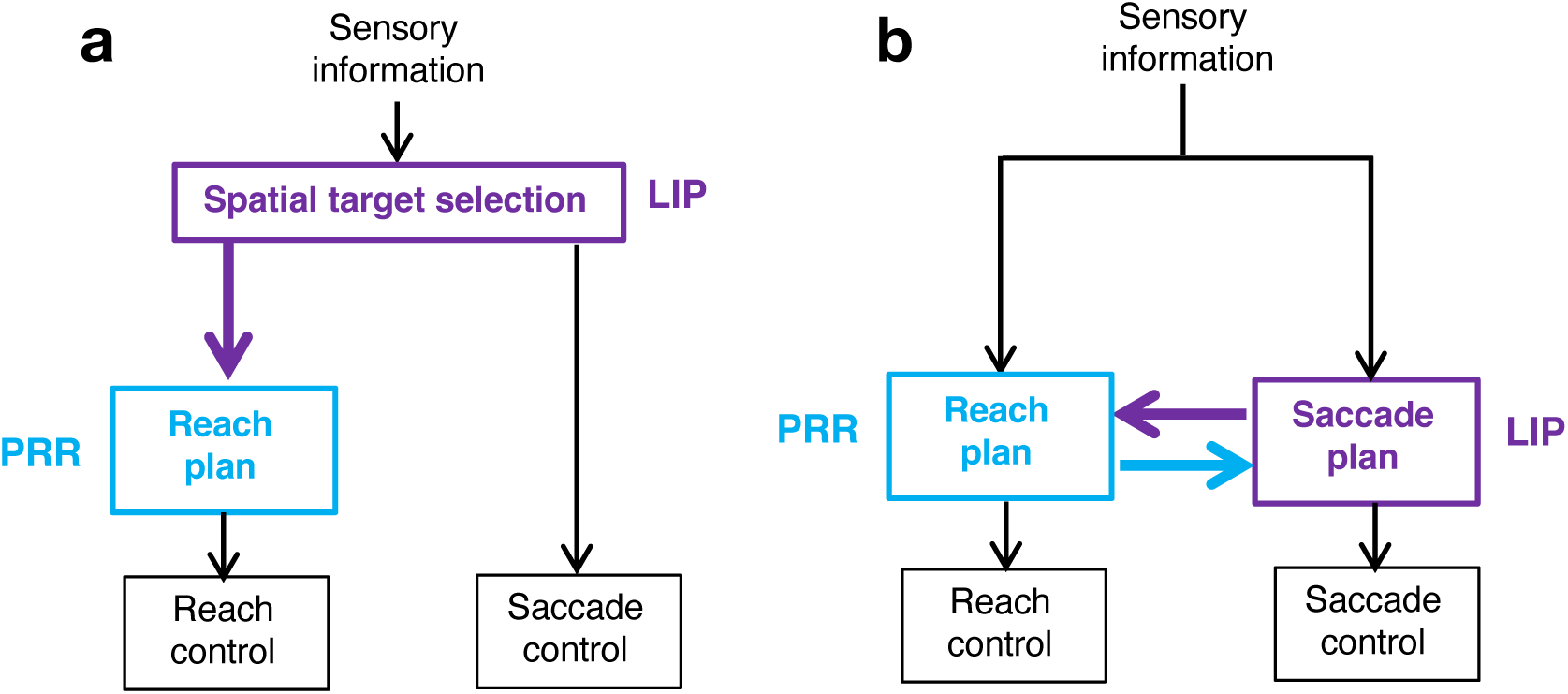
Possible patterns of information flow between PRR and LIP for eye and arm movements. **a**, A hierarchical model. LIP is at a higher hierarchical level than PRR, directing both eye and arm movements. The flow of information is unidirectional: LIP sends spatial target information to PRR (purple arrow) which subserves reach planning, but PRR does not send the same information to LIP. **b,** A parallel model. Sensory information projects to both PRR and LIP. PRR and LIP subserve reach and saccade planning, respectively. The two areas share information in service of eye-hand coordination: PRR sends information about the reach plan to LIP (cyan arrow) and LIP sends information about the saccade plan to PRR (purple arrow).

## Results

We recorded single units and LFP from PRR and LIP in both hemispheres of two monkeys to quantify functional connectivity related to planning eye and arm movements. Single unit activity and LFP power from PRR and LIP are shown in Extended Data Fig. 1. Details of the electrophysiological data are provided in previously published studies^43–46^. Animals reached with the left (unimanual), right (unimanual), or both hands together (bimanual-together) to a single target or moved each hand to a different target (bimanual-apart) (Fig. 2). Animals made saccades to the reach targets on most trials. For single-target trials, a saccade to the target was required, while for bimanual-apart trials animals were free to move their eyes as they chose. In a fifth trial type, animals made a saccade without a reach (saccade-only). All trial types were interleaved. On each trial, animals were instructed to prepare the appropriate movement based on the color of the peripheral target(s) and then cued to initiate that movement after a variable delay period of 1250 to 1750 ms. Overall performance following training was good. 86% of initiated trials (two home buttons touched and initial fixation target acquired) were successfully completed. Saccades slightly preceded unimanual reaches, with median reaction times (RT) of 217 and 328 ms, respectively. Bimanual reaches were initiated roughly synchronously, with a median absolute difference in the two arms’ RTs of 31 and 30 ms, for bimanual-together and bimanual-apart trials, respectively. Additional behavioral details can be found in a previous publication^47^.

**Fig. 2.**
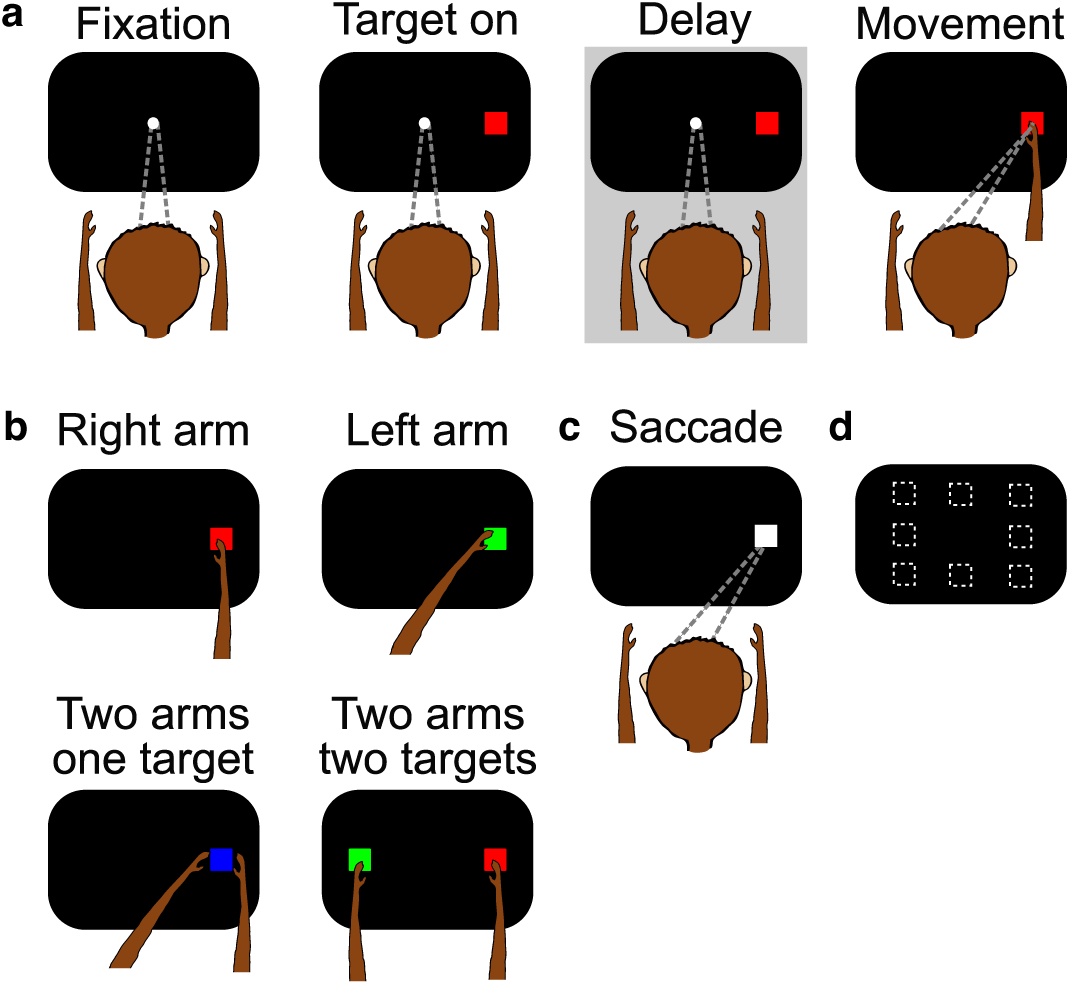
Task design. **A**, Animals begin trials by fixating a central target and placing their hands on home buttons. After 500 ms, a peripheral target appears. The color of the target instructs a particular movement. The target remains on throughout a variable delay period (1250-1750 ms). After the delay, the fixation target disappears, instructing the animal to perform the previously instructed movement (go cue). **b,** A red or green target instructs a right or left unimanual arm movement, respectively; the other hand must stay on its home button. A blue target instructs movements of the two arms to the same target (“bimanual together”). Simultaneous red and green targets instruct movements to separate targets (i.e., right arm to red and left arm to green; “bimanual apart”). Simultaneous targets always appear diametrically opposite to one another. With the exception of bimanual apart trials, the animal is required to make an eye movement to the target at the same time as the arm movement. **c,** Animals also perform saccade-only trials without arm movements. All 5 trial types are interleaved. **d,** Targets can appear at one of eight possible locations. If a RF is identified, only two target locations are used (one in the RF and one diametrically opposed)

### Information flows primarily from PRR to LIP during the movement planning period

We assayed directional information flow between PRR and LIP using two largely independent methods: LFP-LFP spectral Granger causality and time-lagged spike-LFP coherence^16, 48, 49^. We first focus on spike-LFP coherence in the movement planning period, the 800 ms interval prior to the go cue. This interval contains no changes in task-related stimuli that would produce visual responses and no task-related responses that would produce proprioceptive feedback, either of which could drive confounding common input to both PRR and LIP. We use “coherence from A to B” as shorthand for the coherence between spikes in location A with LFP in location B because we expect that spikes in A can more directly drive LFP in B than LFP in A can drive spikes in B (see Discussion). Within a single hemisphere, coherence from PRR to LIP (blue trace) is significantly elevated above chance levels at 10-64 Hz (Fig. 3a). At 25-45 Hz, coherence from PRR to LIP is more than twice as large as coherence from LIP to PRR (purple trace) relative to chance levels (Fig. 3a, pooled t-test, p<0.001 at each frequency). Across hemispheres (i.e., coherence between spikes recorded from one hemisphere and LFP recorded from the other) there is a similar asymmetry, though coherence magnitudes are reduced compared to within-hemisphere effects (Fig. 3b). The same asymmetry can also be found with alternatives to coherence, e.g., pairwise phase consistency, an unbiased measure of spike-field synchronization that is robust to differences in spike counts (Extended Data Fig. 2)^50^.

**Fig. 3.**
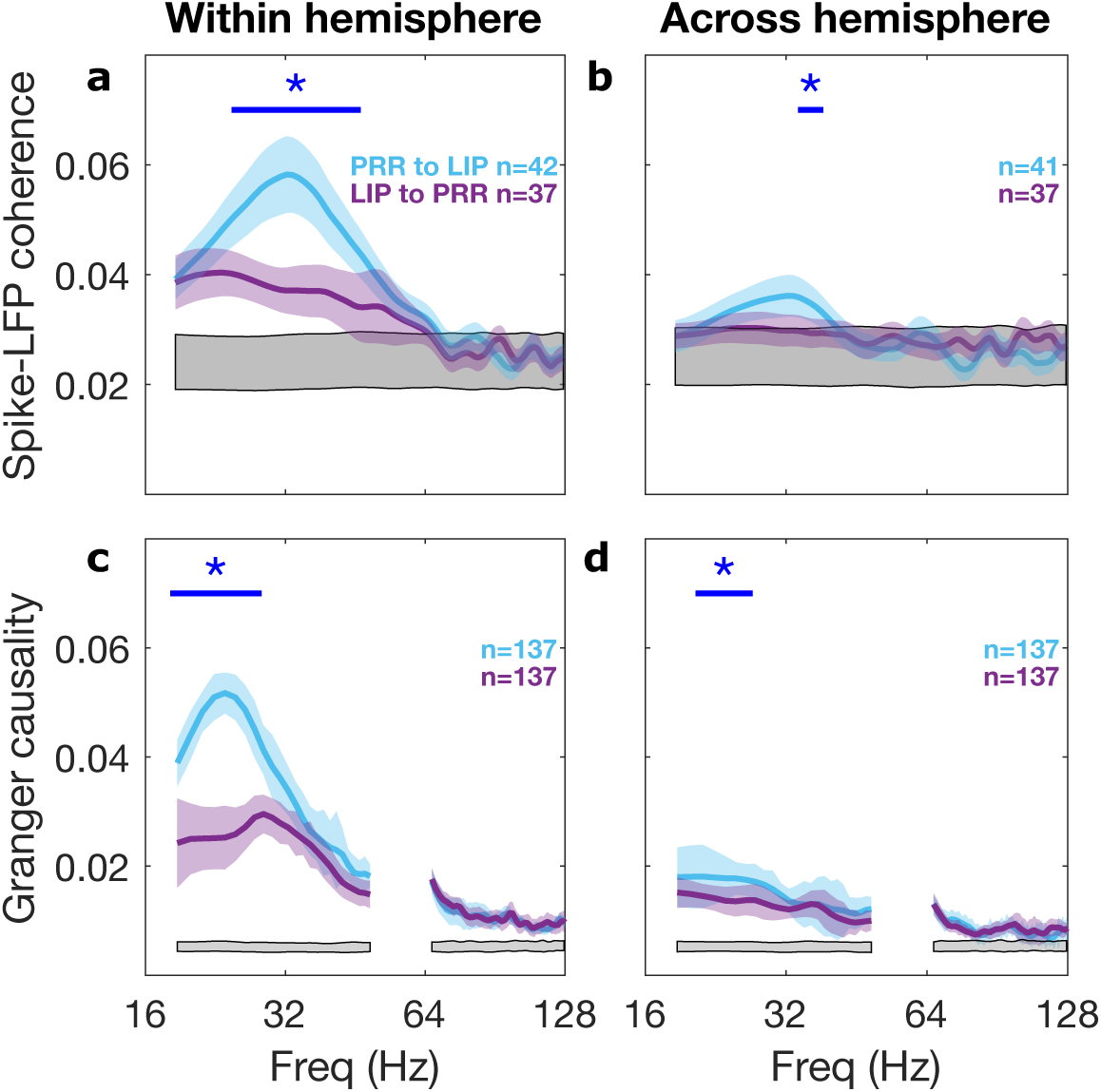
Information flow between PRR and LIP during the planning period for coordinated eye and arm movements. **a**, Within hemisphere spike-LFP coherence is significantly higher from PRR to LIP than vice versa at 25-45 Hz. The blue asterisks and bar denote p<0.001 (pooled t-test). Colored shaded regions denote SEM. The grey shaded region represents the 99% bounds of a shuffle test (See Methods). Peak PRR to LIP coherence is 0.060 at 32 Hz, while peak LIP to PRR coherence is 0.043 at 20 Hz. Measured from the chance level of 0.024, this is a ratio of greater than 2:1. **b,** Spike-LFP coherence across hemispheres is significantly higher from PRR to LIP than vice versa at 35-37 Hz. **c,** Similar effects were found using LFP-LFP Spectral Granger causality. Within hemisphere, peak PRR to LIP Granger causality was 0.052 at 24 Hz, while peak LIP to PRR Granger causality was 0.024 at 30 Hz. The blue asterisks denote p<0.001 (Wilcoxon signed-rank test). Measured from the chance level of 0.005, this is a ratio of 2:1. **d,** Spectral Granger causality between PRR and LIP across hemisphere. PRR to LIP flow is significantly higher than vice versa at 21-26 Hz.

Next, we computed spectral Granger causality between LFPs. Measures of functional connectivity based on LFP-LFP interactions will not necessarily be the same as those based on spike-LFP interactions. However, finding similar directional asymmetries across two independent measures would increase confidence in the results^51^. We use “Granger causality from A to B” as a shorthand for how well LFP from location B can be predicted using LFP from location A. Like spike-LFP coherence, spectral Granger causality is greater from PRR to LIP than from LIP to PRR (within hemisphere, Fig. 3c; across hemispheres, Fig 3d). The directional asymmetry is highly significant at 19-27 Hz (Wilcoxon signed-rank test, p<0.001 at each frequency). Importantly, this directional asymmetry is not due to increased power in one region (Extended Data Fig. 3).

### Information flow from PRR to LIP is task-specific

The data in Fig. 3 were computed after combining responses to all 4 interleaved reach trial types. If PRR transmits information about a planned reach to LIP, then the quality or quantity of information flowing between areas may depend on the type of reach. Indeed, unimanual and bimanual reaches were each associated with unique spike-LFP coherence spectra (Fig. 4a). Within a single hemisphere, the two bimanual tasks were associated with the strongest coherence (0.091) and peaked at 35 Hz. The two unimanual tasks were associated with an intermediate level of coherence and a lower peak (0.087) at 30 Hz. There was significant task-specificity at 27-40 Hz (repeated-measures ANOVA, p<0.001 at each frequency). We observed similar, though not identical, task specificity using different time-frequency bandwidths (data not shown) and with pairwise phase consistency (Extended Data Fig. 4). Cross-hemispheric coherence from PRR to LIP also showed a similar pattern of task-specificity (Extended Data Fig. 5). In contrast, coherence from LIP to PRR was not significantly affected by the type of reach (Fig. 4b). In fact, the coherence from LIP to PRR during reach preparation was barely above the shuffle confidence limits. A similar result is obtained through spectral Granger causality analysis of LFP signals: there is significant task-specific modulation from PRR to LIP at 19-27 Hz but not from LIP to PRR at any frequency (Figs. 4c and d, permutation test, p<0.01).

**Fig. 4.**
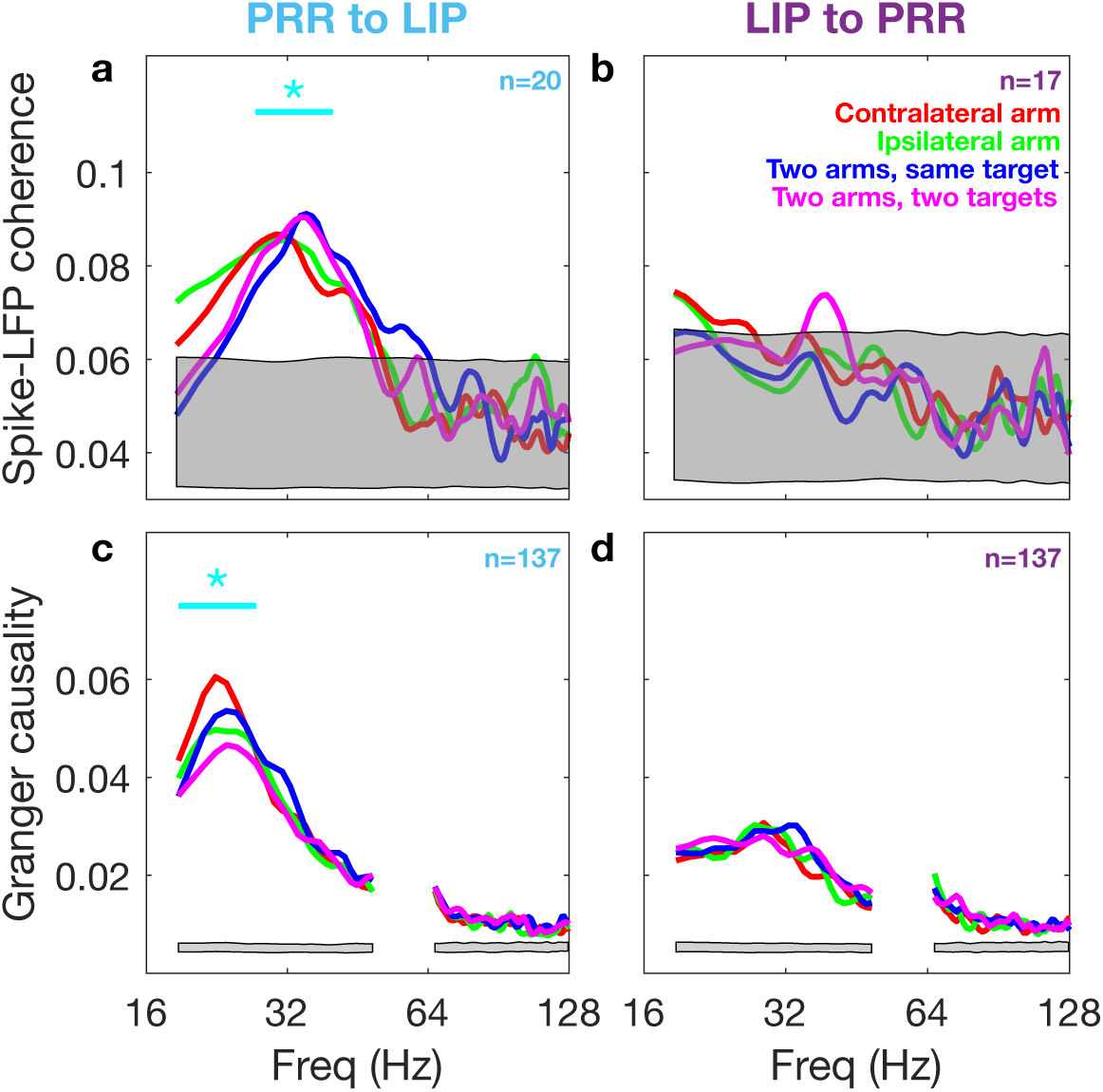
Information flow from PRR to LIP within hemisphere is task-specific. Spike-LFP coherence is shown above (**a-b**) and spectral Granger causality is shown below (**c-d**). **a**, Spike-LFP coherence from PRR to LIP at 27-40 Hz depends on the task type **b,** Spike-LFP coherence from LIP to PRR does not depend on the task type. **c,** Granger causality from PRR to LIP shows task-specific modulation. We tested Granger causality at 19-27 Hz, the frequency range that showed a significant asymmetry (PRR to LIP > LIP to PRR) when all task types were pooled (Fig. 3c). There is significant task-specific modulation at 19-27 Hz. **d**, No task-specific modulation in Granger causality from LIP to PRR. In each panel, grey shaded regions represent the 99% bounds of a shuffle test. Cyan asterisks and straight lines indicate frequencies with a significant variation across reach tasks (repeated-measures ANOVA [p<0.001] for spike-LFP coherence and a permutation test [p<0.01] for spectral Granger causality).

**Fig. 5.**
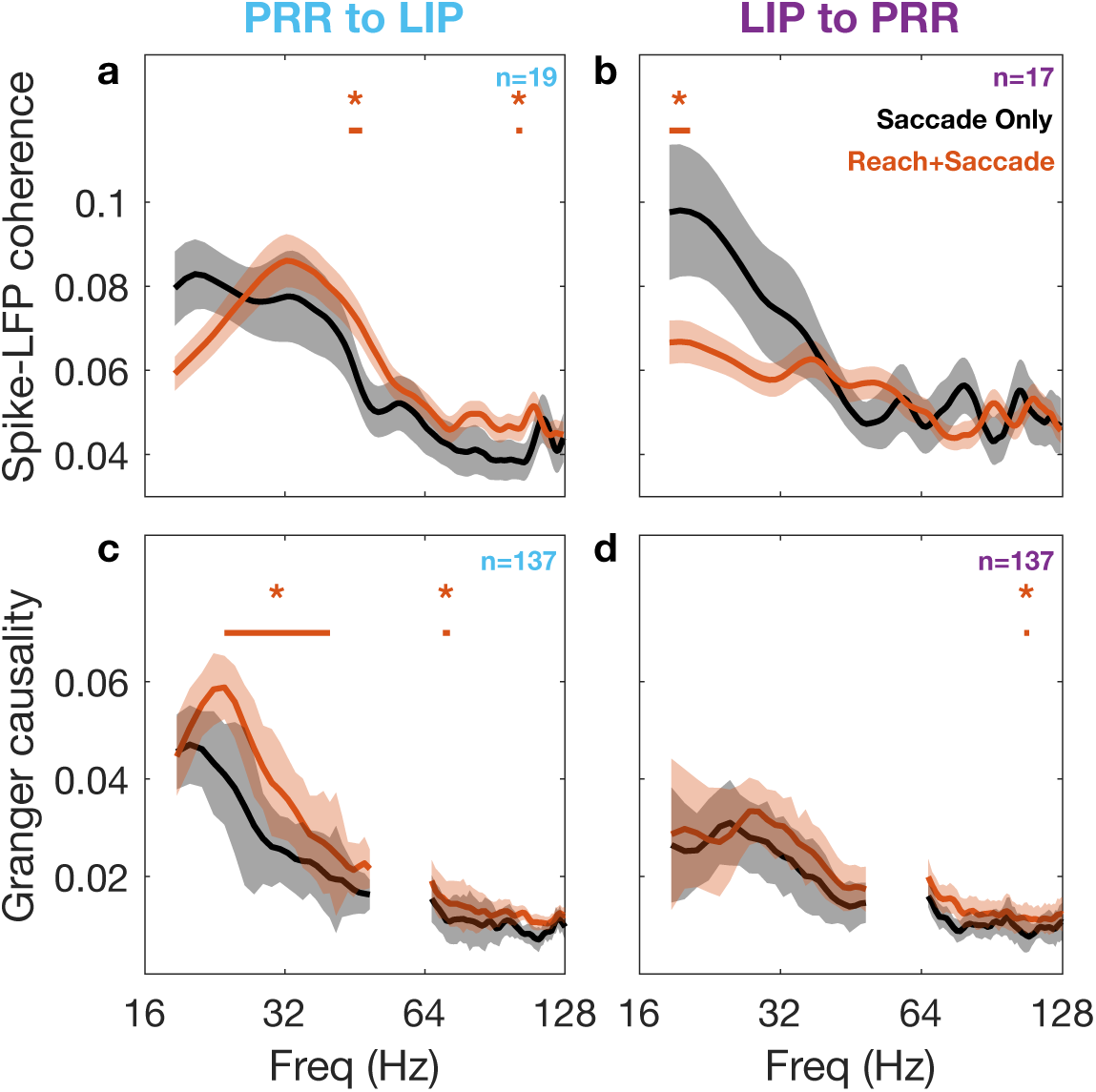
Information flow is effector-specific. Information flow during the preparatory period for coordinated reach plus saccade movements (orange) and for saccade-only movements (black). Spike-LFP coherence is shown above (**a-b**), spectral Granger causality below (**c-d**). Flow from PRR to LIP is shown on the left and from LIP to PRR on the right. **a,** Spike-LFP coherence from PRR to LIP at 0.083 at 21 Hz on saccade-only trials (black) and peaks at 0.086 at 32 Hz on reach plus saccade trials (orange). Coherence for reach plus saccade trials is significantly higher than coherence for saccade-only trials at 44-47 Hz. **b,** Spike-LFP coherence from LIP to PRR peaks at 0.098 at 20 Hz on saccade-only trials (black) and at 0.067 at 20 Hz (orange), respectively. Eye movements (saccade-only) trials have significantly higher coherence than coordinated eye-arm movements at 19-21 Hz. **c,** Granger causality from PRR to LIP has two peaks associated with the two conditions at 0.047 at 20 Hz (black) and 0.059 at 24 Hz (orange), respectively. Coordinated reach plus saccade movements have significantly higher coherence than saccade-only movements at 24-40 Hz. **d,** Granger causality from LIP to PRR shows no effector-specific modulation. In all panels, Orange and black shaded regions denote SEM. Orange asterisks and straight lines indicate frequencies with a significant difference in modulation between the two tasks (Wilcoxon signed-rank test [p<0.01] for both spike-LFP coherence and spectral Granger causality).

We next considered information flow when animals were planning a saccade-only movement compared to a reach plus a saccade (Fig. 5). Peak coherence from PRR to LIP was similar in magnitude for the two conditions but shifted towards higher frequencies for reaches plus saccades compared to saccade-only movements (Saccade: 0.083 at 21 Hz; Unimanual Reach with Saccade: 0.086 at 32 Hz) (Fig. 5a). Coherence from PRR to LIP was higher for planning a reach plus a saccade than a saccade alone at 44-47 Hz (Wilcoxon signed-rank test, p<0.01). In the opposite direction, coherence from LIP to PRR was higher for saccade-only movements than reaches plus saccades at 14-35 Hz (Wilcoxon signed-rank test, p<0.01 at 20 and 22 Hz) (Fig. 5b). Thus, information flow from PRR to LIP was higher when preparing a reach plus a saccade compared to a saccade-only movement while flow from LIP to PRR showed the reverse effect, higher when preparing a saccade-only movement compared to a reach plus a saccade.

Spectral Granger causality between LFP signals also shows effector-specificity, with significantly larger Granger causality values from PRR to LIP for a reach plus a saccade compared to saccade-only at 23-40 Hz, with peak values of 0.059 at 25 Hz versus 0.047 at 20 Hz, respectively (Fig. 5c, Wilcoxon signed-rank test, p<0.01 at each individual frequency). In contrast, Granger causality from LIP to PRR was only half as large and similar for the two tasks: peak values of 0.033 at 28 Hz for a reach plus a saccade and 0.031 at 25 Hz for saccade-only (Fig. 5d).

### Coherence from PRR to LIP reflects a causal effect

An analysis of the temporal lag that maximizes inter-areal spike-LFP coherence can provide information about causal effects. Spikes from one area might drive synaptic and dendritic currents in a second area, thereby contributing to LFP in the second area^52, 53^. Because spike propagation, synaptic transmission, and dendritic conduction are not instantaneous, we expect a lag between when a spike occurs in one area and when that spike exerts its peak influence on LFP in another area (Fig. 6a, d, cyan)^54^. Consider an experiment in which spikes are recorded from area A, LFP is recorded from B, and spike-LFP coherence is computed between these two signals. If information flows from A to B, we expect that spike-LFP coherence would be elevated and would be greatest when the spike signal is lagged (shifted later in time) relative to the LFP signal. If information flows in the opposite direction, from B to A, then we again expect elevated coherence. However, the path for information transfer requires more steps in the case of flow from B to A compared to A to B (compare Figs. 6b and 6a) and so the coherence would be lower for B to A than A to B. Furthermore, coherence for the B to A case should be greatest when the spike signal is moved earlier in time (a negative lag) relative to the LFP signal (Fig. 6d, orange). Finally, elevated coherence could arise from common input to both areas rather than from communication between the areas (Fig. 6c). In this case, the common input will first drive LFP in both areas and then, a short time later, influence spikes in both areas. Thus, coherence will be maximized when a small lead is imposed on the spikes (Fig. 6d, purple).

**Fig. 6.**
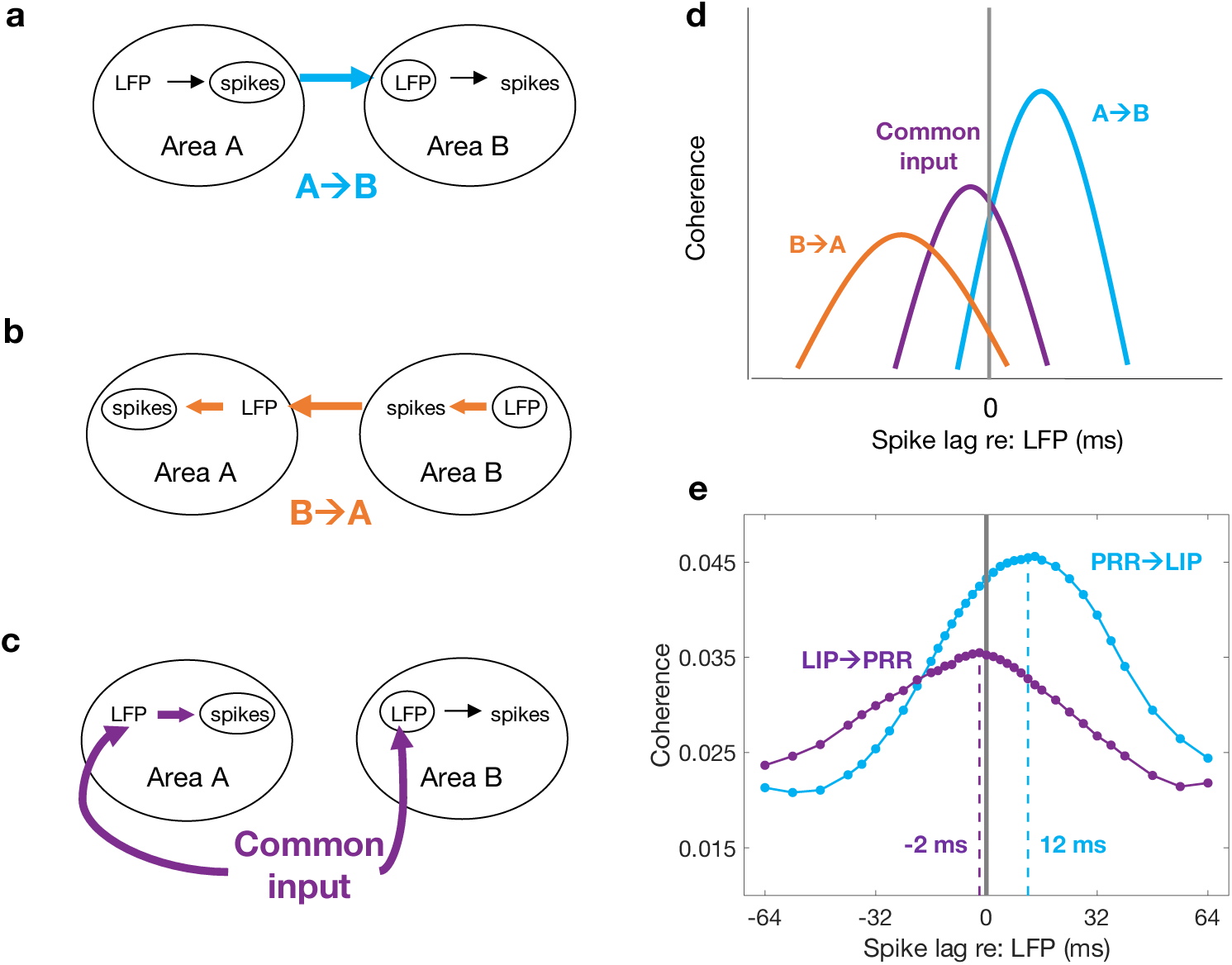
Time-lagged spike-LFP coherence analysis differentiates patterns of information flow. Spikes in area A and LFP in area B (small circles in **a**, **b** and **c**) can share information for a number of reasons. **a,** Information flow from area A to area B (cyan arrow). **b,** Flow from area B to area A (orange arrows). **c,** Common input to areas A and B (purple arrows). **d,** Predicted time-lagged coherence between spikes in area A and LFP in area B for the flows illustrated in panels **a-c**. The X axis indicates whether a lag (X>0) or a lead (X<0) is imposed on spikes from area A relative to LFP in area B prior to computing coherence. With communication from area A to B (cyan), we expect maximal coherence to occur when a lag is imposed on spikes relative to LFP to compensate for conduction times and synaptic delays in the pathway. We expect peak coherence to be strong because spikes in A directly influence LFP in B. With communication from area B to area A (orange), the temporal order is reversed: the LFP in area B (indirectly) influences spikes in area A. As a result, we expect that a lead must be imposed on spikes in area A relative to the LFP in area B in order to maximize coherence. Since the linkage from LFP in area B to spikes in area A is less direct than the previous case, we predict weaker coherence (orange versus cyan). Finally, common input into areas A and B directly influences LFP and indirectly influences spikes in both areas. As a result, we expect inter-areal coherence between spikes and LFP to be maximal when a small lead is imposed on the spikes relative to the LFP (purple). Peak coherence will be smaller than in **a** because the connection is less direct. **e,** Spike-LFP coherence at 25 Hz in all task types from PRR to LIP (cyan) and from LIP to PRR (purple) as a function of spike lag with respect to LFP. The peak lag for PRR to LIP coherence (cyan) at 12 ms is consistent with direct communication (as in **a**). The 2 ms lead and the lower coherence value at peak (0.034 versus 0.044) for LIP to PRR (purple) is consistent with either an indirect effect of PRR to LIP input (as in **b**), common input (as in **c**), or some combination of effects.

We determined the temporal lag or lead that maximizes cross-areal spike-LFP coherence in each direction. From PRR to LIP, there was a clear peak in coherence at 25 Hz when a lag of 12 ms was imposed on spikes relative to LFPs (Fig. 6e, cyan). We repeated this analysis across a broad range of frequencies and found that the spike lag that maximizes spike-LFP coherence (“peak lag”) is significantly different from zero only at 21-36 Hz, with mean values of ∼10 ms across the 5 tasks (Fig. 7a). This timing is consistent with the time required for information to travel from one cortical area to another, and is also consistent with the lag previously reported for high frequency spike-LFP coherence^55–57^. Outside of 21-36 Hz the peak lag was inconsistent (consistent with no transfer of information) or not significantly different from zero. A similar effect was seen for cross-hemisphere interactions (Extended Data Fig. 6). The exact value for peak lag depended on the task being planned, with lags of ∼12 ms when the task included a contralateral reach and smaller values for saccades and ipsilateral reaches (Extended Data Fig. 7a). In the reverse direction, from LIP to PRR, peak coherence occurred with no lag or even a slight lead of ∼4 ms (Fig. 6e, purple; Fig. 7b; and Extended Data Fig. 7b). The overall pattern is consistent with information flowing from PRR to LIP, possibly with the addition of common input into both PRR and LIP. This in turn suggests that PRR, having obtained visual information from earlier visual areas^10^, determines where a reach will be directed and then communicates that information to LIP.

**Fig. 7.**
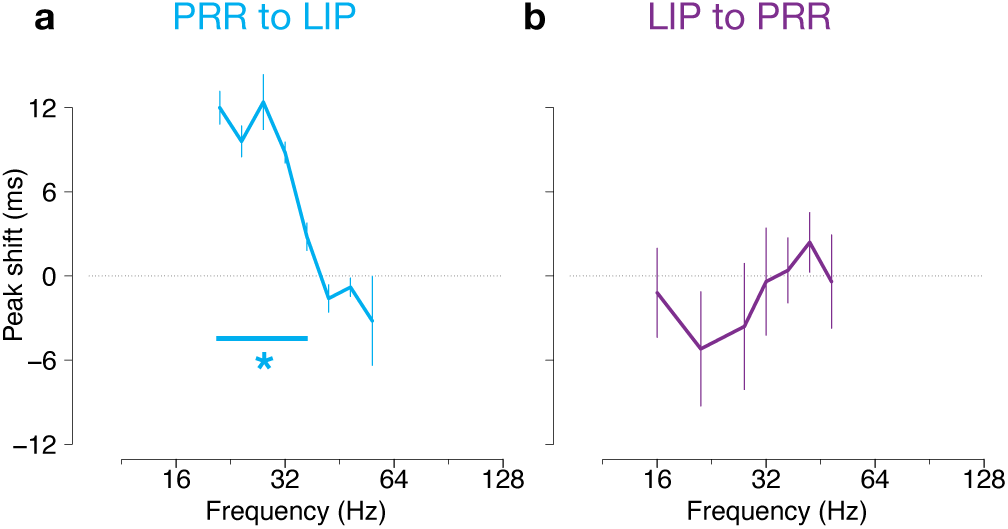
Timed-lagged spike-LFP coherence analysis across different frequencies between PRR and LIP in the same hemisphere. For each direction of communication (from PRR to LIP and from LIP to PRR), we ran the time-lagged analysis for each task type in each frequency and computed an average of peak spike lag (ms) with respect to LFPs from the 5 task types in each frequency. Error bars denote SEM. Data are plotted only for frequencies where the peak lags for all 5 task types were more similar than would be expected by chance (permutation test, p<0.001). **a,** PRR to LIP. At 21-36 Hz, there is a significant ∼10 ms peak lag (two-tailed t-test [p<0.05]), consistent with information flow from PRR to LIP. **b,** LIP to PRR. At 21-36 Hz there is no lag but instead a lead of ∼4 ms (p>0.05), consistent with information flow from PRR to LIP (Fig. 6b).

Our results support the idea that information primarily flows from PRR to LIP during the movement planning period. It is possible, however, that LIP sends information about where to make arm movements to PRR shortly after the target appears, and that the subsequent flow from PRR to LIP reflects feedback to LIP. To test this possibility, we next consider information flow immediately following target presentation.

### Information flow is primarily from PRR to LIP even immediately after target appearance

An alternative interpretation of our data is that LIP sends information to PRR about where to make arm movements after the target appears, and that subsequent information flow, from PRR to LIP, is feedback to LIP. To test this possibility, we next consider information flow immediately following target presentation. In the interval from 50 to 550 ms after target onset, coherence from PRR to LIP was well above chance but coherence from LIP to PRR was substantially smaller and only barely significant (Fig. 8a). The coherence was shifted to slightly higher frequencies than later in the trial, with peaks at 43 and 41 Hz, respectively, but the pattern otherwise resembled the effects during the delay period. There was significant task-specific modulation from PRR to LIP but not vice versa (Extended Data Fig. 8a, b). Looking only at spikes associated with the preferred direction does not change this pattern (Extended Data Fig. 9) and looking at shorter periods starting at target onset (e.g., 50-450 ms) yields noisier but broadly similar results (Extended Data Fig. 10). Granger causality from PRR to LIP and vice versa are both elevated at 20-30 Hz in this early period with no significant difference between the two directions (Fig. 8b). However, there was a significant task-specific modulation in this epoch from PRR to LIP but not from LIP to PRR (Extended Data Fig. 8c, d). Thus, both time-lagged spike-LFP coherence and Granger causality are consistent with PRR playing a command role in specifying spatial target locations both immediately after the target presentation as well as during the preparation period.

**Fig. 8.**
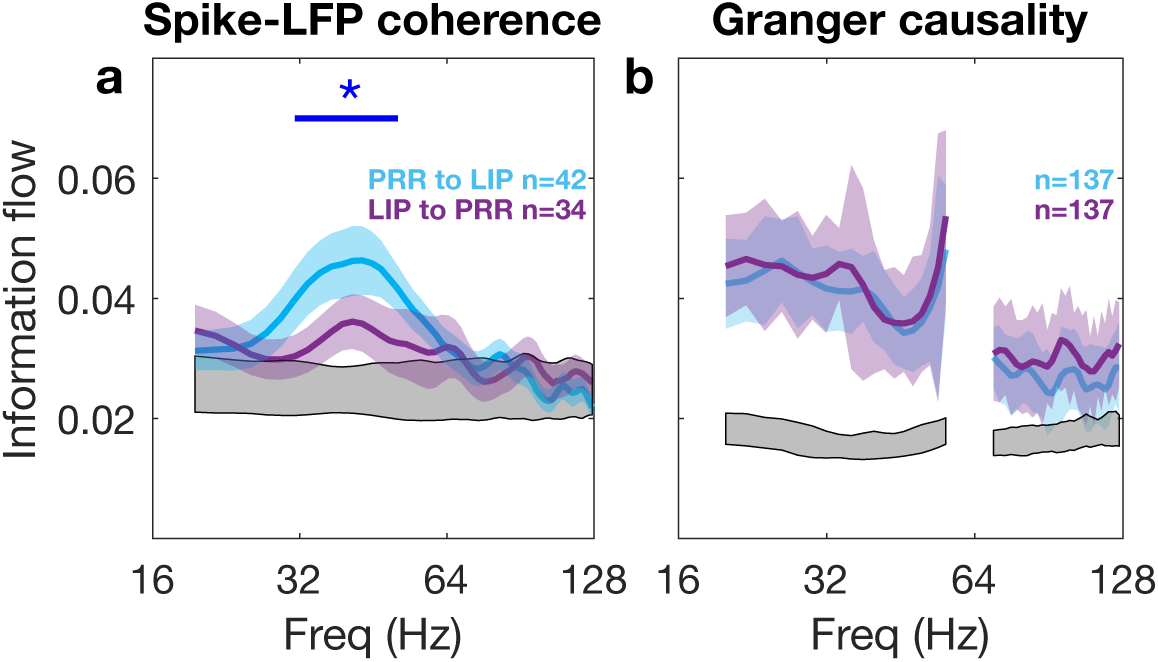
Information flow between PRR and LIP during the target presentation period (50-550 ms aligned to the target onset) for coordinated eye and arm movements. **a**, Spike-LFP coherence between PRR and LIP in the same hemisphere. The blue asterisk and straight line denote p<0.001 (pooled t-test). Peak PRR to LIP coherence was 0.046 at 43 Hz, while peak LIP to PRR coherence was 0.036 at 41 Hz. Measured from the chance level of 0.024, this is a ratio of greater than 2:1. The colored shaded regions denote SEM. The lower grey shaded region represents the 99% bounds from random permutations of information flow from PRR to LIP. **b**, Spectral Granger causality between PRR and LIP in the same hemisphere.

## Discussion

We asked if the relative functional organization of PRR and LIP is best described as hierarchical, with LIP providing high-order spatial information to PRR, or parallel, with each area operating on its own and sharing information on a task-specific basis. To distinguish between these two architectures, we assayed inter-areal information flow across different tasks using time-lagged spike-LFP coherence and spectral Granger causality. In most of the tasks we tested, information flow from PRR to LIP was twice as large as the flow from LIP to PRR (Fig. 3a, c). Flow from PRR to LIP was higher for behaviors involving reaches compared to saccades alone (Fig. 5a) and depended on the particular type of reach being performed (Fig. 4a, c). Flow from LIP to PRR was higher for saccades alone compared to behaviors involving reaches (Fig. 5b) and did not depend on the type of reach being performed (Fig. 4b, d). This is consistent with a parallel rather than hierarchical architecture for the planning of reaching and saccades.

Previous work has shown that PRR represents targets for planned arm movements while LIP represents targets for planned eye movements^22–26^. Eye movements are tightly and reciprocally coupled to attention: we look at what we attend to, and our attention automatically shifts to the goal of an upcoming saccade^58^. This reciprocal relationship is evident during the planning period prior to an eye movement, even in peripheral attention tasks in which an eye movement is disallowed^59^. Thus it is not surprising that LIP is active not just with saccades but also in non-saccadic tasks involving spatial attention. On the basis of such activations, it has been proposed that LIP forms a “priority map” of space^35, 36^ which could then propagate behaviorally-relevant spatial information to other areas to guide not just saccades but other movements as well^34, 60–62^. Coordinated eye and arm movements provide an excellent test of this proposal. Primates, including humans, often look where they will reach, and saccade and reach RTs are correlated^63–67^. It is known that LIP also contains information about planned arm movements^60, 68, 69^. It is not clear, however, whether LIP specifies the reach target or if the decision about where to reach is made elsewhere, for example in PRR, and then propagated to LIP to coordinate eye and arm movements. A dominant role for LIP in reaching is suggested by the fact that visual responses appear sooner in LIP than PRR and by the fact that saccades usually precede reaches^70^. However, other observations suggest that PRR and LIP might operate in a parallel and reciprocal manner. Reach preparation enhances visual processing^71, 72^, and saccades coupled with reaches are executed more quickly than singular saccades^67^. Further, saccades and reaches can be spatially decoupled, and when decoupled, attention can be allocated separately to each effector’s target^73–76^.

Our main finding is that, during the planning period prior to a reach, information flows primarily from PRR to LIP (Fig. 3). If LIP was a command center for spatial information processing, then brain regions such as PRR would be consumers of that information and information would primarily flow from LIP to PRR. Instead, we find that information flow is bidirectional, with twice the flow from PRR to LIP compared to LIP to PRR. Our approach relies on identifying similarities in the signals contained within PRR and LIP and then asking whether one signal is likely to be driving the other. There is a long history of using Granger causality analysis for this purpose, asking if the past history of one signal can predict the current modulation of the second signal, after taking into account the past history of the second signal^77^. Spectral Granger causality analysis is a variant thereof that operates in frequency space rather than in the time domain but follows similar principles^78, 79^. In order to increase confidence in our results we used a second independent method of assessing information flow: time-lagged spike-LFP coherence^16^. Action potentials (spikes) propagate information along axons. Spikes give rise to synaptic currents, which in turn give rise to dendritic currents. These currents are thought to produce much of the modulation of the LFP^52, 53^. Spike-LFP interactions have an inherent directional asymmetry. Spikes in area A can directly evoke an LFP response in area B (Fig. 6a), but the reverse direction requires three steps: LFP in area B reflects synaptic and dendritic currents that can drive spikes in area B, these spikes can drive currents in area A, and these currents can drive spikes in area A (Fig. 6b). This asymmetry means that coherence between spikes in area A and LFP in area B is more likely to reflect a causal influence from area A to area B (a single step) rather than from area B back to area A (three steps). We tested this idea by computing coherence between spikes in one area and LFP in another after time-shifting them by different amounts, and then asking whether peak coherence occurred when a lag was imposed on spikes relative to LFPs (consistent with a causal influence from the area in which the spikes were recorded to the area in which the LFP was recorded) or when a lead was imposed on spikes (consistent with either common input to both areas or information flow from the area in which the LFP was recorded to the area in which the spikes were recorded). Our results confirm that spikes in PRR drive LFP in LIP: peak coherence from PRR to LIP occurred when a lag of ∼10ms was imposed on spikes relative to the LFPs (Fig. 6e, cyan). This is long compared to axonal propagation and synaptic conduction (a few milliseconds) but consistent with the average difference in response latencies for direct connections between cortical areas and with previous reports of the timing of activity across areas^55–57^. Taken together, the LFP-LFP spectral Granger causality, the spike-LFP coherence analyses, and the findings from the time-lagged spike-LFP analysis, provide high confidence for the conclusion that more information flows from PRR to LIP than from LIP to PRR.

An alternate interpretation of our results that preserves a dominant role of LIP in guiding reaches is that LIP sends spatial information to PRR early in the trial, shortly after the target first appears. In this view, the information flow that we observe during the planning period is merely feedback, perhaps used to ensure that PRR has received the correct information from LIP. We rule out this possibility by showing that, in the first 300, 400 or 500 ms of the trial, information still flows predominantly from PRR to LIP, not from LIP to PRR (Fig. 8 and Extended Data Fig. 10). Only during a saccade-only task do we see higher information flow from LIP to PRR, associated with lower temporal frequencies than during reach trials (Fig. 5).

A pitfall of correlation-based analyses is that communication may reflect common input from a third area rather than direct communication between areas. To minimize this possibility, we focused on the movement preparation period when no new stimuli were presented and no task-related actions were performed, either of which might drive robust activity that might then serve as common input into PRR and LIP. The time-lagged spike-LFP coherence analysis rules out common input as the main driver for flow from PRR to LIP (spikes in PRR, LFP in LIP; Figs. 6 and 7). With common input, coherence would be maximized when a lead was imposed on the spikes relative to the LFPs. Instead, peak coherence occurs when a lag is imposed on the spikes.

A second pitfall of this study is our implicit assumption that information flow, measured by lagged coherence, is monotonically related to causal influence. Information flow from PRR to LIP was twice that from LIP to PRR. This suggests that PRR has more causal influence on LIP than vice versa, but this assumes that the information being encoded and the efficiency of the encoding is similar for transfer in the two different directions. Finally, our coherence approach captures only linear relationships between the activity of the two regions. More sophisticated mathematical techniques such as generalized linear models or transfer entropy could be used to capture additional non-linear relationships^80, 81^.

In summary, we quantified functional connectivity between PRR and LIP using time-lagged spike-LFP coherence and spectral LFP-LFP Granger causality to investigate the functional organization of micro-circuits in posterior parietal cortex. PRR and LIP encode plans for arm and eye movements, respectively, but LIP has also been implicated in high level abstract spatial processing. The former suggests parallel positions in cortical processing streams, while the latter suggests a more hierarchical organization, with LIP specifying spatial targets to PRR and similar regions. We found that the pattern of causal influences between the two areas supports the conclusion that PRR plays a command role in spatial target selection and that the two areas operate in parallel, with PRR determining reach targets and LIP determining saccade targets.

## Online Methods

### Apparatus

Experiments took place in a dark room. Head-fixed animals sat in a custom-designed monkey chair (Crist Instrument, Hagerstown, Maryland) with an open front to allow unimpaired reaching movements with both arms. Visual stimuli were back-projected by an LCD projector onto a translucent plexiglass screen mounted vertically ∼40 cm in front of the animal. Eight target positions on the screen were organized in a rectangle centered on the fixation point, each target ∼8 cm (11°) or ∼11 cm (15°) from the center fixation point. At each target location, a small piece of plexiglass (5 cm × 1 cm) oriented in the sagittal plane was mounted on the front of the projection screen to bisect the touching surface. The animals were trained to reach with the left and right hands to the left and right sides, respectively, of the plexiglass divider. Touches were monitored every 2 ms using 9 pairs of capacitive sensors. One pair of sensors served as home pads to detect reach starting points. Each of the remaining sensor pairs were place behind a target position, one on each side of the plexiglass, to detect reach endpoints. Thus for every target, each hand activated a unique capacitive sensor, even when both hands reached to the same target. Eye position was monitored using an infrared video eye-tracking system (120 Hz ISCAN eye-tracking laboratory, ETL-400).

### Behavioral tasks

The task design and the movement conditions are shown in Fig. 2. The animals performed delayed saccade-only movements or coordinated eye and arm movements with the left, right, or both arms. Animals first fixated on a circular white stimulus (1.5° x 1.5°) centered on the screen in front of them. Left and right hands touched home pads situated at waist height and 20 cm in front of each shoulder. After holding fixation (± 5°) and initial arm positions for a fixed duration of 500 ms, either one or two peripheral targets (5° × 5°) appeared on the screen for 1250-1750 ms. Fixation was required throughout this instructed delay period. After the delay, the central eye fixation target shrank in size to a single pixel, cueing the animal to move to the peripheral target(s) in accordance with target color. A blue target instructed a reach with both arms (“bimanual-together”). A green or red target instructed a reach with the left or right arm, respectively. The simultaneous appearance of two targets (red and green) cued a reach with both arms to two different targets (“bimanual-apart”). Only trials in which the two targets were separated by 180° relative to the central fixation point are used for the current report (i.e., reaches to the left and right, top and bottom, or opposed diagonal locations). For bimanual-apart reaches, the arms could be uncrossed or crossed. Finally, a white target instructed a saccade without a reach. To help ensure as natural coordination as possible, animals were not trained to make arm movements without accompanying eye movements. All single-target reach trials require an accompanying saccade to the target. Saccades were optional (but almost always performed) for two-target reach trials.

All trial types were randomly interleaved within sets of 10 or 40 trials (one each per condition [5] and direction [2 or 8]). Throughout saccade and unimanual reach trials, hands not instructed to move were required to remain on their respective home buttons. On bimanual trials, the left and right hands were required to hit their target(s) within 500 ms of one another. Animals were required to maintain their hand(s) on the final target(s) for 300 ms. Spatial tolerance for saccades was ±5°. When an error occurred (a failure to achieve or maintain the required eye or hand positions), the trial was aborted and short time-out ensued: 1500ms for an early fixation break and 500ms for a targeting error. Aborted trials were excluded from further analyses. Successful trials were rewarded with a drop of water or juice.

### Electrophysiological recordings

Recordings were made from the left and right hemispheres of two adult male rhesus monkeys. In each animal, two recording chambers were centered at ∼8 mm posterior to the ear canals and ∼12 mm lateral of the midline on each side and placed flush to the skull. Anatomical magnetic resonance images were used to localize the medial bank of the intraparietal sulcus. Extracellular recordings were made using glass-coated tungsten electrodes (Alpha Omega, Alpharetta, GA; electrode impedance 0.5-3.0 MΩ at 1kHz) recorded from a steel guide tube in the same recording well. Neural signals were processed and saved using the Plexon MAP system (Plexon, Inc.). Signals were passed through a pre-amplifier and then separated into two signal paths. The LFP channel was band-pass filtered between 0.7 to 300 Hz and digitized at 1 kHz. We used a band-pass filter to remove 60 Hz power from the LFP. The spike channel was band-pass filtered between 100 Hz and 8 kHz and digitized at 25 kHz. Single units were isolated online via manually-set waveform triggers. During each recording session, one or two electrodes were placed in PRR and LIP in each hemisphere, up to a total of 4 electrodes. While searching for cells, animals performed saccade-only trials and combined reach plus saccade trials with the contralateral arm (contralateral with respect to the side of the isolated cell). Online, the preferred direction for a cell was defined as the target location that resulted in the largest sustained firing during the delay period. The null direction was defined as the diametrically opposite direction. Data were then collected for all trial types. As an indirect marker of inter-areal communication, we computed both inter-areal spike-LFP coherence and LFP-LFP spectral Granger causality over a broad range of frequencies.

### Spike-LFP coherence

Coherence spectra between spikes in one area and LFP signals in another were computed over the last 800 ms before the go cue and from 50 to 550 ms after target onset. Because spike-LFP coherence is biased towards higher values for smaller numbers of spikes, we only included spike-LFP pairs with at least 500 spikes when combining over task types (e.g. Figs. 3a,b, 9a, Extended Data Figs. 9, 10a) or 300-400 spikes when computing coherence separately for each task type (e.g., Figs. 4a,b, 7a,b, Extended Data Figs. 5a,b, 8a,b). Key findings were confirmed using pairwise phase consistency, an alternative method of computing coherence that circumvents this bias^50^. We computed spike-LFP coherence using a multitaper method implemented in the Chronux toolbox (http://chronux.org/)82. The number of tapers was typically 9 (but see next section).

Mean coherence spectra were estimated as follows. First, Fourier transforms were computed for each trial, *n*, and each taper, *k*, according to Eqs. (1) and (2) with *t* as the the sample index, *T* as the the number of samples per time series, *f* as the frequency in Hz, *j* as the imaginary unit 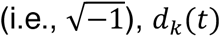 as the taper time series for taper *k*, and *x_n_*(*t*) and *y_n_*(*t*) as the spike or LFP time series for trial *n*. For the spike time series, the DC component, i.e., the average value over time, was subtracted before transformation.

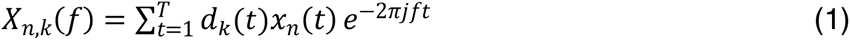

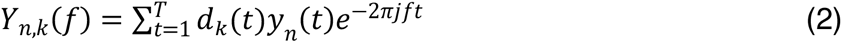

The power spectral densities for a single trial, *S*_*xx*,_(*f*) and *S*_*yy*,*n*_(*f*), were then computed as an average of the cross-spectra across *K* tapers according to Eqs. (3) and (4).

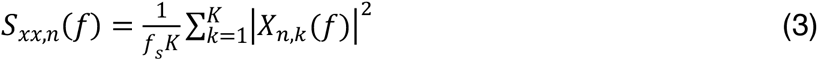

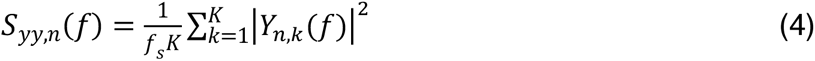

*f_s_* is the sampling frequency, and *K* is the number of tapers. The power spectral densities, *S*_*x*_(*f*) and *S*_*yy*_(*f*), were then averaged across *N* trials to produce a single estimate of the power spectral density according to Eqs. (5) and (6).

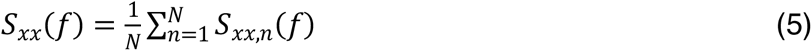

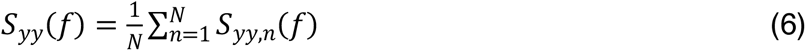

Next, a mean cross power spectrum, *S_x,y_*(*f*), was computed by averaging spectral estimates across *K* tapers and *N* trials as in Eqs. (7) and (8) where 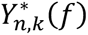 represents the complex conjugate of *Y_n,k_*(*f*).

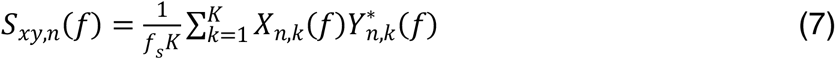

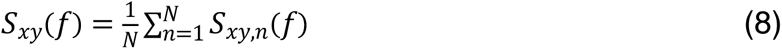

Coherence was computed by normalizing the cross spectrum by the geometric mean of the power spectra as in Eq. (9). Finally, coherence was averaged across spike-LFP pairs; note that coherence and thus its average are complex valued.

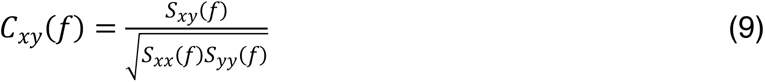

### Time-lagged spike-LFP coherence analysis

To further investigate inter-areal signal flow, we asked if there is a clear lag or lead in spikes relative to LFP that maximizes spike-LFP coherence. A lag in spikes is consistent with direct communication (Fig. 6a), and a lead is consistent with either communication in the reverse direction or common input (Fig. 6b, c). We calculated coherence after imposing a lag or lead on the spikes with respect to the LFPs from -128 ms to +128 ms. We then asked, for a given LFP frequency, what lag or lead maximized coherence (peak shift). A representative example of the analysis centered at 25 Hz is shown in Fig. 6e. We repeated this at 20 different logarithmically-spaced frequencies from 9 to 128 Hz. A pitfall of this analysis is that even if there is no shared information, there will always be some lag that produces maximum coherence. In that case, the peak shift would be equally likely to occur anywhere between -128 and +128 ms. For the main analysis (Fig. 7), peak shifts were computed separately for each of the 5 tasks, using 23 tapers and a frequency half-bandwidth of 15 Hz. The data were then plotted only for those center frequencies in which the peak shifts for the 5 tasks were clustered. Statistically significant clustering was determined using a randomization test and a criterion value of p<0.001.

### Spike-LFP pairwise phase consistency (PPC)

The LFP phase at the time of each spike was estimated with a wavelet transform but was not pooled across trials. PPC was estimated according to Eq. (10) (Extended Data Figs. 2, 4).

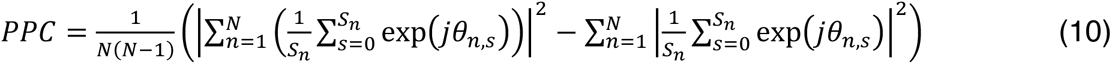

*N* is the total number of trials, and *S_n_* is the number of spikes in trial *n*, and *θ_n,s_* is the phase at the time of spike *s* in trial *n*.

### Spectral LFP-LFP Granger causality

We recorded 137 pairs of LFPs in PRR and LIP in the same hemisphere (56 and 81 pairs in each animal, respectively), and 137 pairs in different hemispheres (58 and 79 pairs in each animal, respectively). We used nonparametric bivariate spectral Granger causality to quantify causal relationships between these signal pairs using the FieldTrip toolbox^48^. Granger causality assays information flow from signal *x* to signal *y* by quantifying how much of the variance of signal *y* can be explained by the recent history of the two signals together compared to just the history of signal *y* alone^77^. If this difference is large, then signal *x* is presumed to causally influence signal *y*. Results from time-based autoregressive models, however, depend on the model order and may not fully take spectral characteristics of data into account^83, 84^. To study frequency-specific causal interactions between LFP signals, we calculated spectral Granger causality *GC_x_*_→_*_y_*(*ω*) from a signal *x* to another signal *y* at frequency *ω* to estimate causal interactions using cross-spectral density matrix factorization according to

Eqs. (11) and (12)^78, 79^.

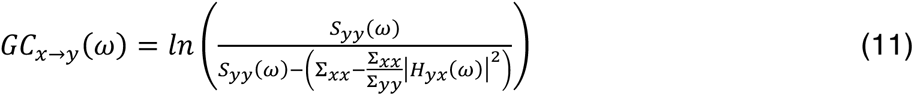

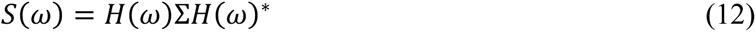

*S*(*ω*) and *H*(*ω*) are the cross-spectral density matrix and the spectral transfer matrix for a pair of signals at frequency *ω*, respectively. Σ is the covariance of an autoregressive model’s residuals. Note that effects below 16 Hz were unreliable (Extended Data Fig. 3).

### Statistics

All trial types were randomly interleaved for each cell or site in a recording session. Statistical analyses were performed in MATLAB (Mathworks) and R Statistical Software (version 4.2.0; The R Foundation for Statistical Computing). All statistical tests were two-sided unless specified otherwise. Chance levels of coherence and Granger causality were computed using permutation analysis (1000 and 200 repetitions, respectively). For spike-LFP coherence and PPC, spike times were randomized within each trial by permuting the interspike intervals. For Granger causality, LFP signals from each electrode in a pair were permuted across trials. Gray regions show the 1st to 99th percentiles of the permuted trials (Figs. 3, 4, 8, and Extended Data Figs. 2, 4, 5, 8, 9, 10); values above these gray regions are significant at the p<0.01 (one-tailed) level. To test significance for between pairs of conditions (Figs. 3, 8 and Extended Data Figs. 2, 3, 9, 10), pooled t-tests or Wilcoxon signed-rank tests were applied at each frequency. Because Granger causality is a biased statistic with a minimum of 0 but no upper bound, we used Wilcoxon signed-rank tests for Granger causality values. Computing the appropriate multiple comparisons correction for the ∼80 tests is difficult because data values at nearby frequencies are not independent, and frequency smoothing in the analysis exacerbates these dependencies. We therefore set a conservative criterion of p<0.001 at three or more contiguous frequency bands and at least one condition to be outside of the 1st and 99th percentile bounds of the permutation tests. When considering the saccade task in isolation (Fig. 5), we have only 1/5 to 1/4 as many trials as in other analyses that involve all trials or all reach trials. To compensate for the resulting loss of power, we relaxed the criterion to 3 or more consecutive points at p<0.01 rather than p<0.001.

## Acknowledgements

This work was supported by the National Eye Institute Grant EY-012135 (L.H.S.) and the Washington University Cognitive Computational and Systems Neuroscience Fellowship (J.K.). We would like to thank Drs. Charles Holmes and David Kaplan for their comments on the manuscript.

## Author contributions

Experimental design: E.M. and L.H.S.

Data collection: E.M.

Data analysis: J.K. and L.H.S.

Original manuscript: J.K. and L.H.S.

Manuscript editing: J.K., E.M., and L.H.S.

## Data availability

All the relevant data are available upon reasonable request. Inquiries should be directed to the corresponding author.

## Code availability

All the relevant codes are available upon reasonable request.

---

**Extended Data Fig. 1.**
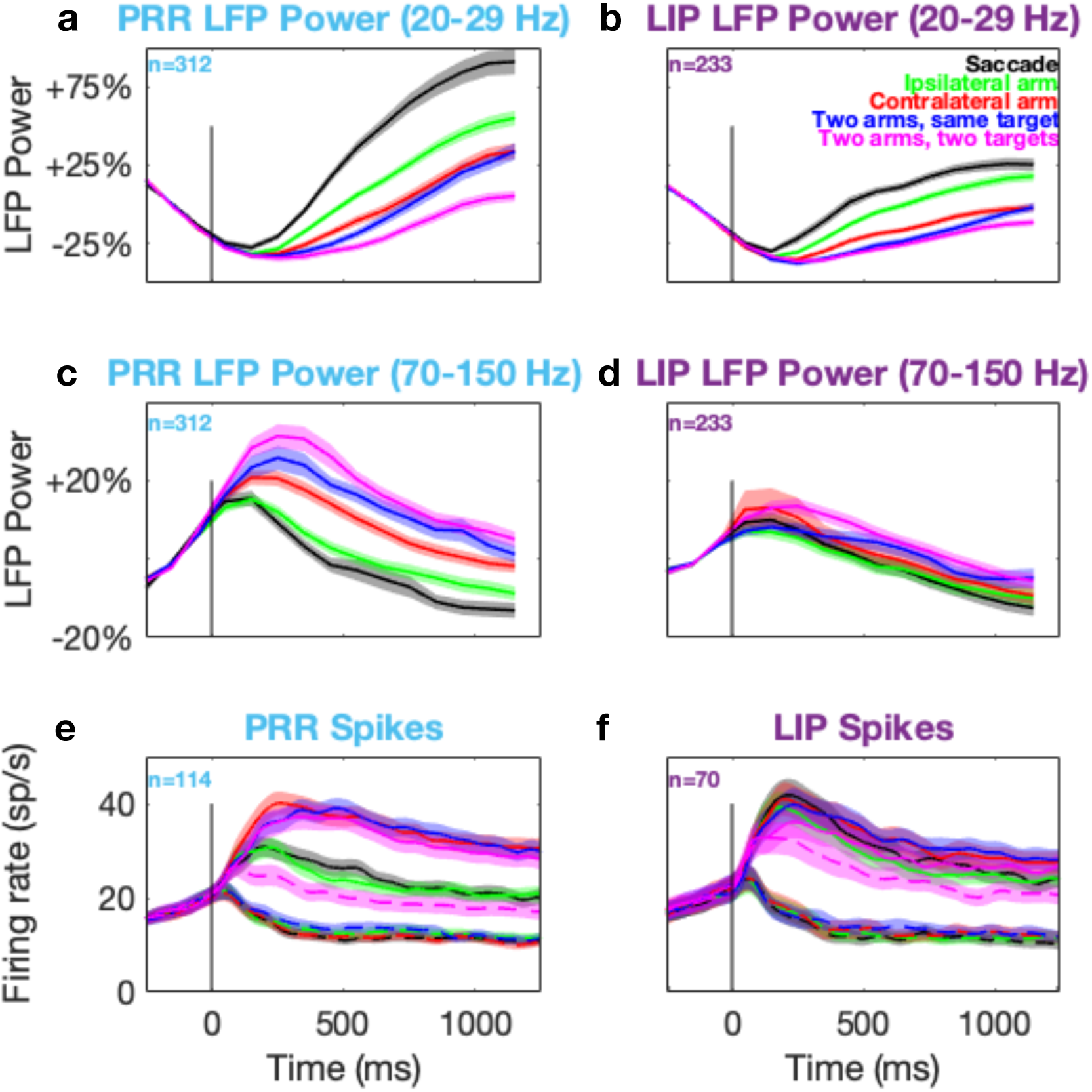
Spikes and LFP power from PRR and LIP aligned to target onset. **a**, LFP power in PRR at 20-29 Hz. **b,** LFP power in LIP at 20-29 Hz. **c,** LFP power in PRR at 70-150 Hz. **d,** LFP power in LIP at 70-150 Hz. **e,** Spikes in PRR. **f,** Spikes in LIP. Solid lines in all panels denote preferred direction. Shaded regions in all panels denote SEM. Dotted lines in (**e-f**) denote null direction.

**Extended Data Fig. 2.**
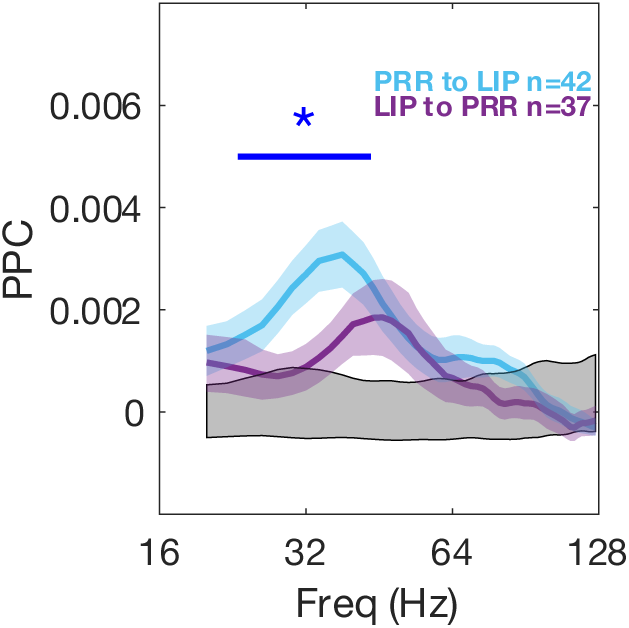
Spike-LFP pairwise phase consistency (PPC) between PRR and LIP within hemisphere. Format identical to Fig. 2a. For both directions of communication (from PRR to LIP and from LIP to PRR), we computed spike-LFP PPC after combining the four reaching tasks during the planning period. PRR to LIP PPC is significantly higher than vice versa at 24-42 Hz. Peak PRR to LIP PPC is 0.031 at 38 Hz, while peak LIP to PRR PPC is 0.019 at 46 Hz. Measured from the chance level of 0.000, this is a ratio of greater than 1.6:1. The grey shaded regions represent the 99% bounds of a shuffle test. Colored shaded regions denote SEM. The blue asterisk and straight line denote p<0.001 (pooled t-test).

**Extended Data Fig. 3.**
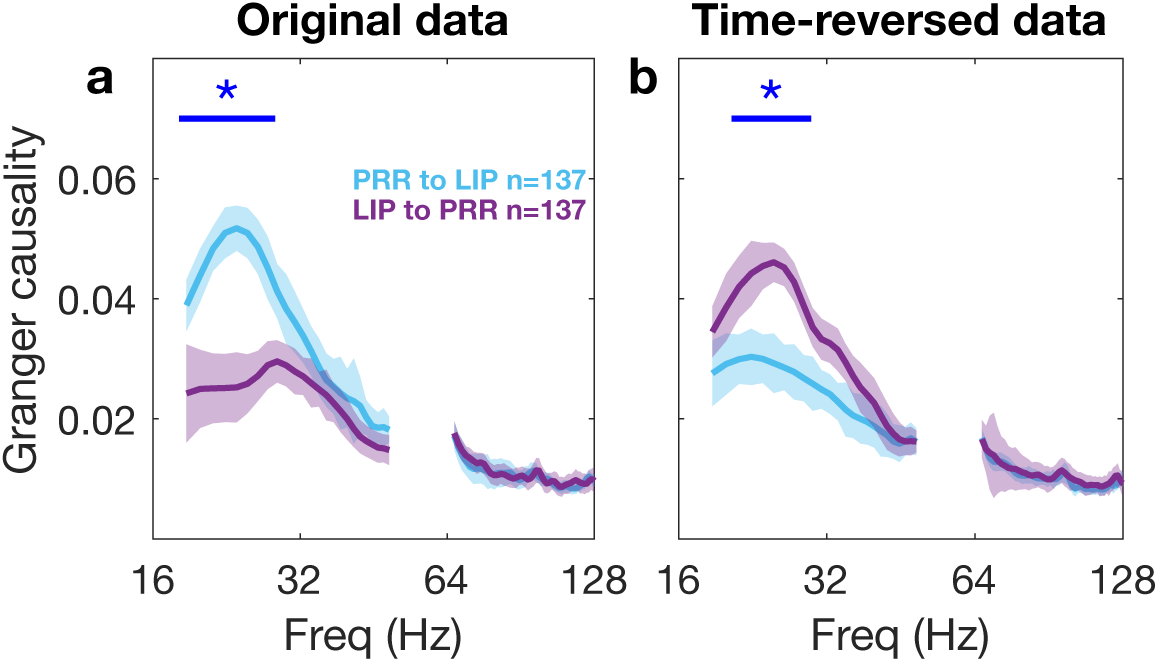
Time-reversal analysis of spectral Granger causality between PRR and LIP. Time reversal should invert the order of the two traces. If this is not the case, then Granger causality results are likely due to an artifact such as greater power in one signal than the other^56–58^. In both panels, colored shaded regions denote SEM. The blue asterisks and straight lines denote p<0.001 (Wilcoxon signed-rank test). **a,** Granger causality from original data in during the planning period of coordinated eye and arm movements, 800 ms prior to the cue to initiate movement. Granger causality from PRR to LIP is significantly higher than Granger causality from LIP to PRR at 19-27 Hz. **b,** Granger causality from time-reversed data. With time-reversal, Granger causality from PRR to LIP is significantly lower than Granger causality from LIP to PRR at 21 to 29 Hz. Peak PRR to LIP Granger causality was 0.030 at 22 Hz, while peak LIP to PRR Granger causality was 0.046 at 25 Hz. This suggests that the observed effect is legitimate rather than due to artifact. In contrast, effects below 16 Hz were not inverted (data not shown), indicating that Granger causality from these low frequencies is unreliable.

**Extended Data Fig. 4.**
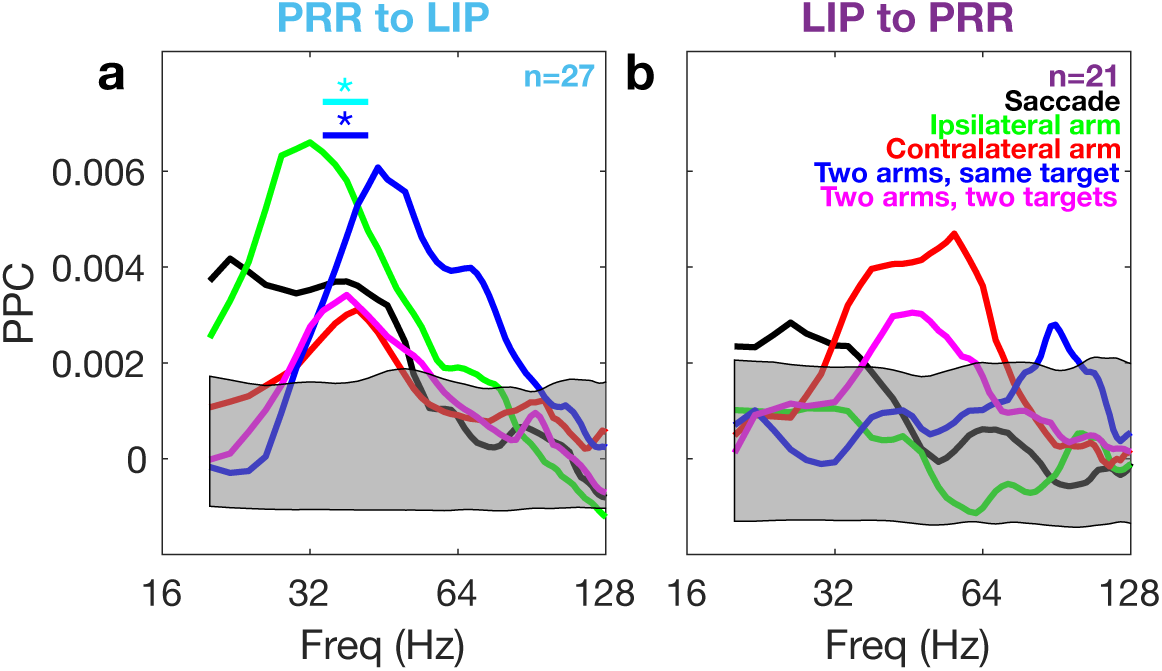
Spike-LFP pairwise phase consistency (PPC) is task-specific from PRR to LIP. For each direction of communication (from PRR to LIP and from LIP to PRR, respectively), we compute spike-LFP PPC for each task type at each frequency during the planning period of coordinated eye and arm movements. The grey shaded regions represent the 99% bounds of a shuffle test. **a,** From PRR to LIP. Blue and cyan asterisks and straight lines indicate frequencies (34-42 Hz) with significant differences in PPC modulation in all five tasks and the four tasks that involve reaching, respectively (repeated-measures ANOVA [p<0.01]). **b,** From LIP to PRR. No significant task-specific modulation in PPC.

**Extended Data Fig. 5.**
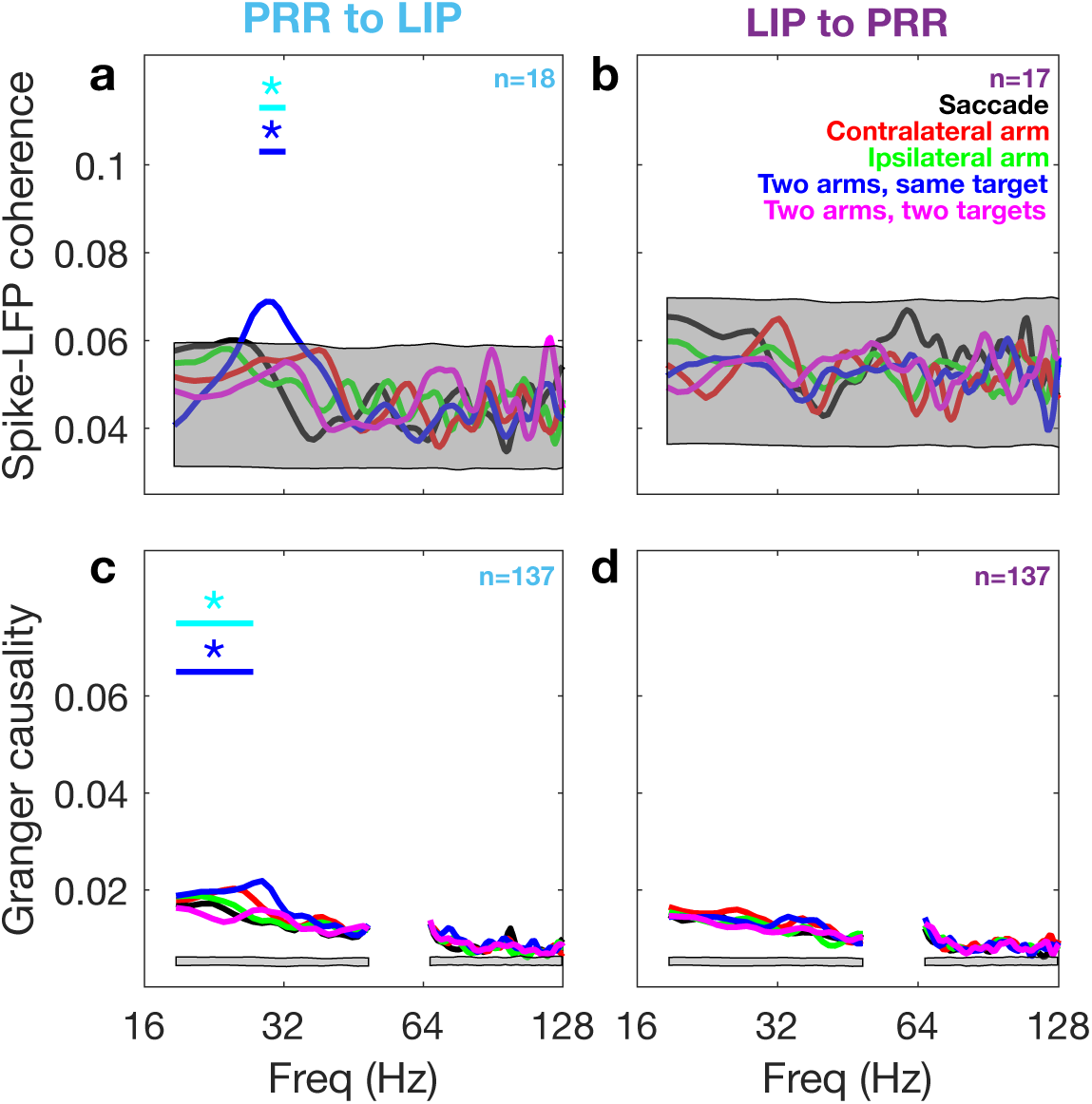
Information flow is task-specific from PRR to LIP across hemispheres. Spike-LFP coherence is shown above (**a-b**), spectral Granger causality below (**c-d**). The grey shaded regions represent the 99% bounds of a shuffle test. Cyan and blue asterisks and straight lines indicate frequencies with a significant difference in modulation across four reach tasks and all five tasks, respectively (repeated-measures ANOVA [p<0.001] for spike-LFP coherence, and a permutation test [p<0.01] for spectral Granger causality). **a,** Spike-LFP coherence from PRR to LIP. The effects are not strong; only the two arms, same target task (blue) has a peak at 0.069 at 29 Hz above the 99% upper bound of the grey shaded region. However, there is a significant task-specific modulation at 28-32 Hz. **b,** Spike-LFP coherence from LIP to PRR. All spike-LFP coherence values are within the 99% bounds of the grey shaded region. No task-specific modulation. **c,** Granger causality from PRR to LIP. There is task-specific modulation at 19-27 Hz. Peak values in Granger causality from PRR to LIP are at 20-30 Hz: saccade (black) at 0.017 at 22 Hz, contralateral arm (red) at 0.020 at 25 Hz. ipsilateral arm at 0.019 at 20 Hz, two arms, same target (blue) at 0.022 at 29 Hz, and two arms, different targets at (magenta) at 0.016 at 19 Hz. **d,** No task-specific modulation in Granger causality from LIP to PRR.

**Extended Data Fig. 6.**
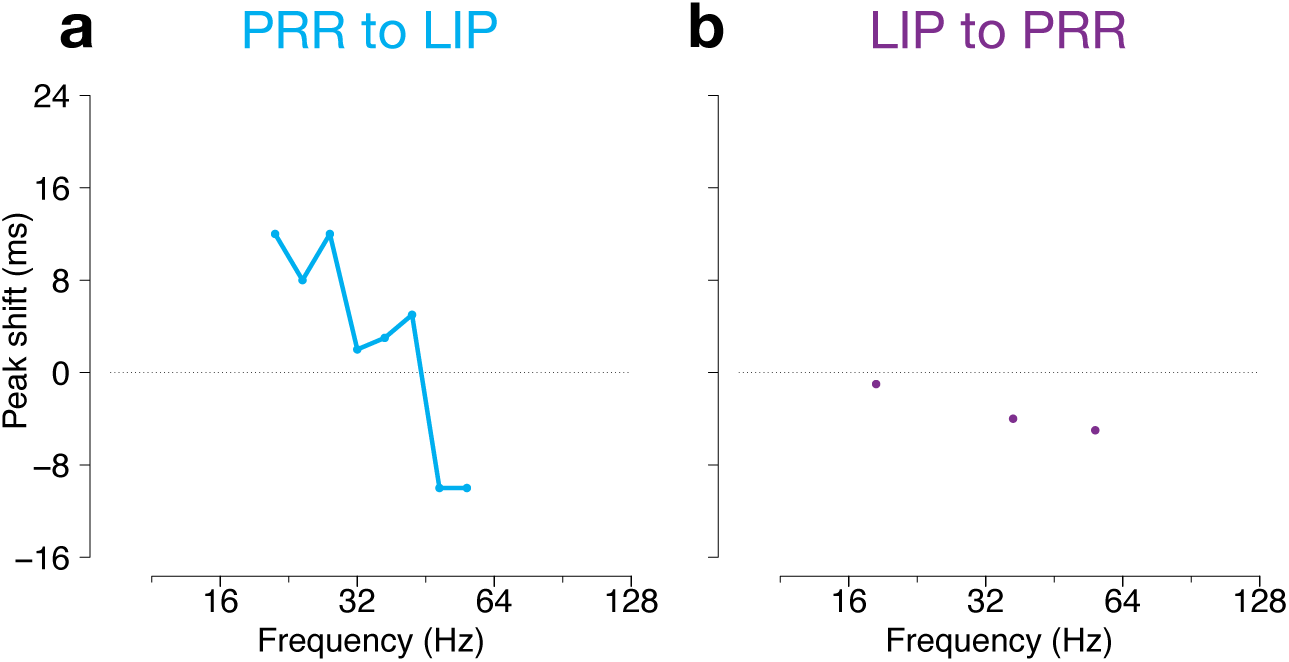
Timed-lagged spike-LFP coherence analysis across different frequencies between PRR and LIP in different hemispheres. For each direction of communication (from PRR to LIP and from LIP to PRR in different hemispheres), we ran the time-lagged analysis for each task type in each frequency and computed a peak spikes lag (ms) with respect to LFPs by merging data in all of the five task types in each frequency. a, From PRR to LIP. At 21-36 Hz, there is a clear peak of ∼10 ms lag. b, From LIP to PRR. At most frequency ranges, there was no clear peak or lag. At 21-36 Hz, there was no single frequency data point where a task type had a significant peak or lag.

**Extended Data Fig. 7.**
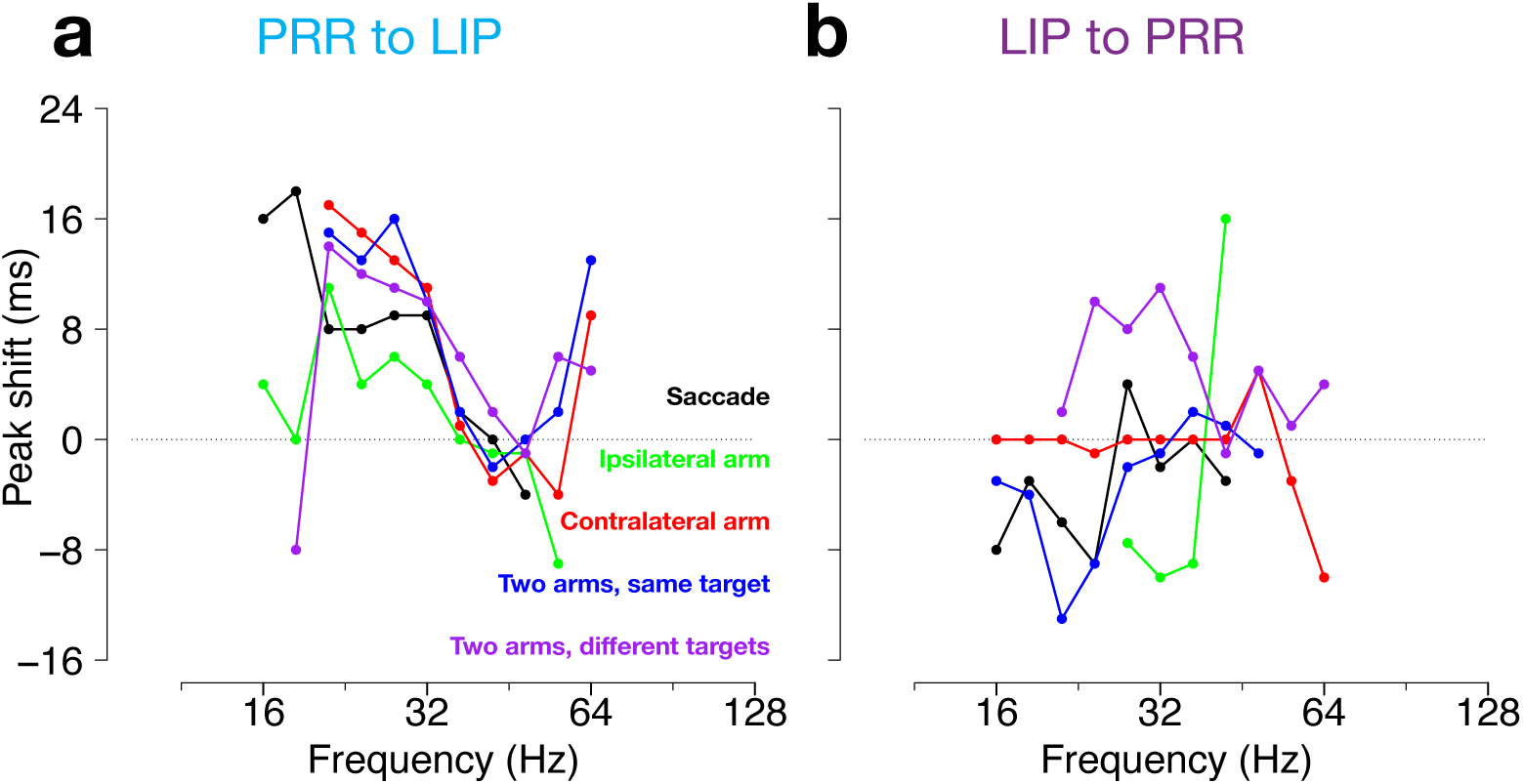
Timed-lagged spike-LFP coherence analysis across different frequencies in each task. For each direction of communication (from PRR to LIP and from LIP to PRR within hemisphere, respectively), we ran the time-lagged spike-LFP coherence analysis for each task type in each frequency and computed a peak time shift (ms) with respect to LFPs. Frequencies with no clear peak are omitted. **a,** From PRR to LIP. At 21-36 Hz, saccade (black) and ipsilateral reach (green) have lower peaks of ∼8 ms. The other task types which include contralateral reach (red, blue, purple) have peaks of ∼12 ms. **b,** From LIP to PRR. At 21-36 Hz, bimanual movements to different targets (purple) have a peak of ∼8 ms and contralateral reach (red) has no consistent lag or lead. The other task types (black, blue, green) have a lead of ∼8 ms, which is consistent with common input.

**Extended Data Fig. 8.**
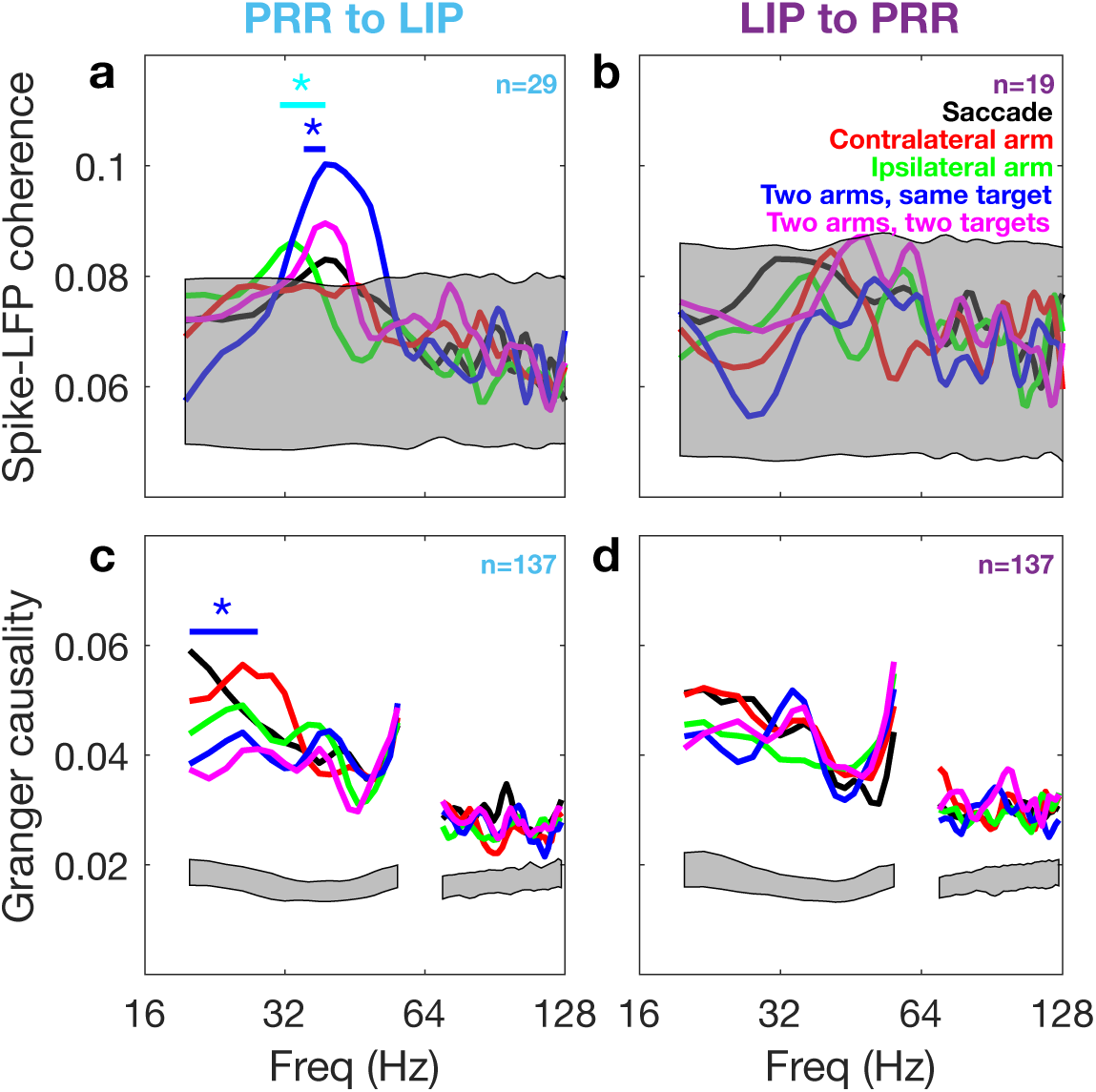
Information flow 50-550 ms after target onset on coordinated eye and arm movement trials (colored) and eye movement only trials (black). Spike-LFP coherence is shown above (a-b), spectral Granger causality below (c-d). Flow from PRR to LIP is shown on the left and from LIP to PRR on the right. The lower grey shaded region represents the 99% bound of a shuffle test. Cyan and blue asterisks and straight lines indicate frequencies with a significant difference in modulation by four reach tasks and all five tasks, respectively (repeated-measures ANOVA [p<0.001] for spike-LFP coherence, and a permutation test [p<0.01] for spectral Granger causality). a, Spike-LFP coherence from PRR to LIP. Peak coherence values were at 39 Hz: 0.083 (saccade, black), 0.100 (two arms, same target, blue), and 0.090 (two arms, different targets, magenta). Ipsilateral arm movements (green) have a peak at 0.086 at 33 Hz. There is a significant task-specific modulation across four reach task types and all five task types at 31-39 Hz (cyan asterisk) and 35-39 Hz (blue asterisk), respectively. b, No task-specific modulation in spike-LFP coherence from LIP to PRR. All coherence values are within the 99% bounds of the grey shaded area. c, Granger causality from PRR to LIP. Granger causality from PRR to LIP has significant task-specific modulation at 19-27 Hz across four reach task types (p<0.01, blue asterisk) and all five task types (p<0.02). Peak coherence for saccade-only (black) is at 0.059 at 20 Hz. All four reach task types have their respective peak coherence values at 26 Hz: 0.056 (red), 0.049 (green), 0.053 (blue), and 0.041 (magenta). d, No task-specific modulation in Granger causality from LIP to PRR. Bimanual movement task types (blue and magenta) have their peaks at 0.052 at 34 Hz and 0.948 at 36 Hz, respectively. Saccade and unimanual movement task types have their respective peaks at 22 Hz: 0.052 (black and red) and 0.046 (green).

**Extended Data Fig. 9.**
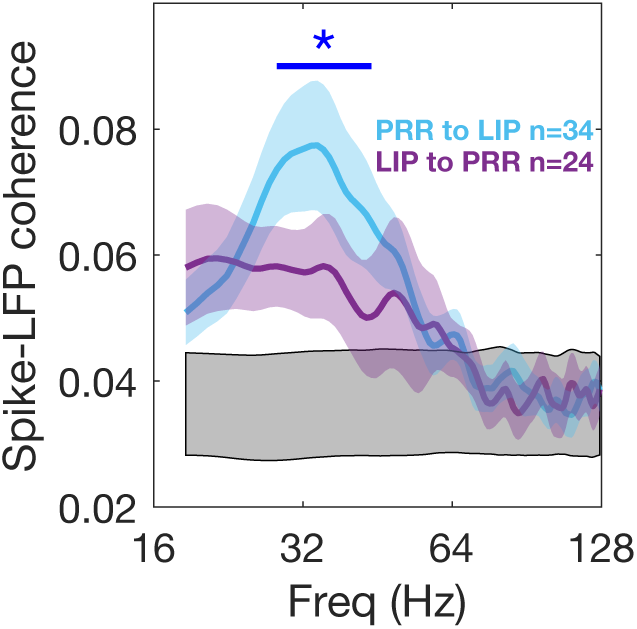
Coherence between spikes associated with the preferred direction and LFP between PRR and LIP within hemisphere. PRR to LIP coherence is significantly higher than vice versa at 28-44 Hz. The blue asterisk and straight line denote p<0.001 (pooled t-test). Peak PRR to LIP coherence is 0.077 at 34 Hz, while peak LIP to PRR coherence is 0.060 at 21 Hz. Measured from the chance level of 0.036, this is a ratio of greater than 1.7:1. Colored shaded regions denote SEM. The lower grey shaded region represents the 99% bounds of a shuffle test.

**Extended Data Fig. 10.**
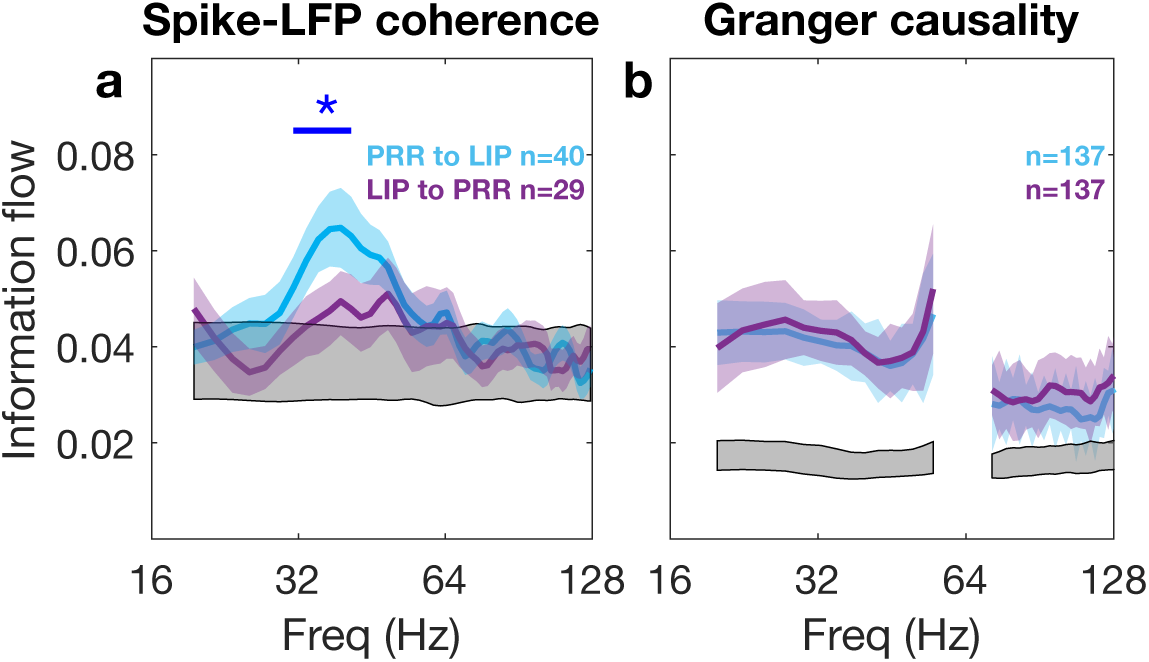
Information flow between PRR and LIP during the target presentation period (50:450 ms aligned to the target onset) for coordinated eye and arm movements. **a**, Spike-LFP coherence between PRR and LIP in the same hemisphere. Peak PRR to LIP coherence was 0.065 at 39 Hz, while peak LIP to PRR coherence was 0.050 at 39 Hz. Measured from the chance level of 0.036, this is a ratio of greater than 2:1. The colored shaded regions denote SEM. The lower grey shaded region represents the 99% bounds from a shuffle test. The blue asterisk and straight line denote p<0.001 (pooled t-test). Granger causality from PRR to LIP was significantly higher than vice versa at 31 to 42 Hz. In a shorter target presentation period (50:350 ms aligned to the target onset), spike-LFP coherence from PRR to LIP was higher than vice versa at 27 to 37 Hz (pooled t-test [p<0.01]). **b**, Spectral Granger causality between PRR and LIP in the same hemisphere. No significant differences between information flow between the two directions.

